# Ras/MAPK modifier loci revealed by eQTL in *C. elegans*

**DOI:** 10.1101/127274

**Authors:** Mark G. Sterken, Linda van Bemmelen van der Plaat, Joost A. G. Riksen, Miriam Rodriguez, Tobias Schmid, Alex Hajnal, Jan E. Kammenga, Basten L. Snoek

**Author notes:** The raw transcriptomic data is available at ArrayExpress (E-MTAB-5856). Corresponding authors: Jan Kammenga, Laboratory of Nematology, Wageningen University, Droevendaalsesteeg 1, 6708PB, Wageningen, The Netherlands.; Basten Snoek, Laboratory of Nematology, Wageningen University, Droevendaalsesteeg 1, 6708PB, Wageningen, The Netherlands.

## Abstract

The oncogenic Ras/MAPK pathway is evolutionarily conserved across metazoans. Yet, almost all our knowledge on this pathway comes from studies using single genetic backgrounds, whereas mutational effects can be highly background dependent. Therefore, we lack insight in the interplay between genetic backgrounds and the Ras/MAPK-signaling pathway. Here, we used a *Caenorhabditis elegans* RIL population containing a gain-of-function mutation in the Ras/MAPK pathway gene *let-60* and measured how gene expression regulation is affected by this mutation. We mapped eQTL and found that the majority (~73%) of the 1516 detected *cis*-eQTL were not specific for the *let-60* mutation, whereas most (~76%) of the 898 detected *trans*-eQTL were associated with the *let-60* mutation. We detected 6 eQTL *trans*-bands specific for the interaction between the genetic background and the mutation, one of which co-localized with the polymorphic Ras/MAPK modifier *amx-2*. Comparison between transgenic lines expressing allelic variants of *amx-2* showed the involvement of *amx-2* in 79% of the *trans*-eQTL for genes mapping to this *trans*-band. Together, our results have revealed loci hidden loci affecting Ras/MAPK signaling using sensitized backgrounds in *C. elegans*. These loci harbor putative polymorphic modifier genes that would not have been detected using mutant screens in single genetic backgrounds.

## INTRODUCTION

The Ras/MAPK pathway is highly conserved across metazoans and regulates a wide range of physiological responses, such as cell proliferation, apoptosis, cell differentiation, and tissue morphogenesis (Gokhale and Shingleton 2015). In humans, activating (“gain-of-function”) mutations in HRas and KRas are strong tumor initiating mutations (Prior and Hancock 2012). Activation of MAP kinase components in model organisms has been shown to cause cell transformation and is implicated in tumorigenesis (Mansour et al. 1994; Cowley et al. 1994). As a key pathway in vertebrates and invertebrates, Ras/MAPK-signaling has been thoroughly studied in model organisms. Genetic studies in the model nematode *Caenorhabditis elegans* have provided insight into *let-60* Ras/MAPK signaling. Activated LET-60, a member of the GTP-binding RAS family (Beitel et al. 1990; Han and Sternberg 1990), induces the phosphorylation of LIN-45 (a Raf ortholog), MEK-2 (a MAPK kinase), and MPK-1 (an ERK ortholog) (Wu and Han 1994). After phosphorylation, MPK-1 enters the nucleus where it regulates many genes by phosphorylation of transcription factors (Tan et al. 1998). Additionally, *let-60* activation underlies programmed cell death in *C. elegans* (Jiang and Wu 2014).

In *C. elegans* almost all studies on *let-60* activation have been conducted with mutant screens using single genetic backgrounds (*i.e*. a mutation in one genotype, in this case Bristol N2). However, the phenotype of induced mutations can vary widely depending on the genetic background (Duveau and Felix 2012; Chandler et al. 2014; Schmid et al. 2015; Kammenga 2017). Induced mutations in one genetic background do not reveal the allelic effects that segregate in natural populations and contribute to phenotypic variation (Kammenga et al. 2008). At the moment we lack insight into how genetic background effects modulate activated Ras/MAPK signaling and which genetic modifiers are involved in the underlying genetic architecture of gene expression.

Here we go beyond mutant screens of *let-60(gf)* in a single genetic background in *C. elegans* by incorporating the effects of multiple genetic backgrounds. This provides the opportunity to explore the genetic variation for identifying novel modifiers of the Ras/MAPK pathway. We recently mapped modifiers affecting Ras/MAPK signaling associated with vulval development in a population of recombinant inbred lines derived from a cross between wildtype Hawaiian CB4856 and Bristol N2 (Schmid et al. 2015). The lines were sensitized by introgression of the G13E gain-of-function mutation (*n1046)* in the Ras gene *let-60* (mutation introgression recombinant inbred lines, miRILs). This mutation is analogous to mutations causing excess cell division in human tumors (Kyriakakis et al. 2015). Hawaiian CB4856 males were crossed with Bristol N2 *let-60* mutants. Random segregation of the two parental genomes was allowed, except for the *let-60* mutation which was kept homozygous from the F2 generation onwards. After ten generations of self-fertilization, to drive all regions to homozygosity, independent miRILs were successfully obtained, each carrying the mutation. Quantitative trait locus (QTL) analysis on the vulval index (VI) of these miRILs in combination with allele swapping experiments revealed the polymorphic monoamine oxidase A (MAOA) gene *amx-2* as a negative regulator of Ras/MAPK signaling (Schmid et al. 2015).

Here, we extended the study by Schmid *et al*. by studying the transcriptional architecture of the same *let-60(gf)* sensitized miRILs. First, we measured the transcriptome of 33 miRILs using microarrays. Second, to gain further insight in the underlying molecular mechanisms, pathways, and new modifiers we mapped gene expression QTL (eQTL). eQTL are polymorphic loci underlying variation in gene transcript abundances and can be used to detect loci which affect many transcript levels and transcriptional pathways (Jansen and Nap 2001). We found that the *let-60(gf)* mutation mainly reveals novel *trans*-eQTL, which are eQTL distant from the regulated gene. In contrast, *cis*-eQTL – genes that co-localize with the eQTL – were mostly independent of the *let-60(gf)* mutation. Of the *trans*-eQTL, 77 genes have an eQTL in a *trans*-band (eQTL hotspot) co-localizing with *amx-2*. Comparing the transcriptional profiles of transgenic lines with the N2 *amx-2* allele with the CB4856 *amx-2* allele showed the involvement of *amx-2* in 79% (61/77) of the genes mapping to this locus. Through network-assisted gene expression analysis, we found evidence that *amx-2* indirectly affects gene expression mapping to this *trans*-band.

## MATERIAL AND METHODS

### General methods and strains used

The strains MT2124 (N2 background with the *let60(n1046)* gain of function mutation) and CB4856 were used, as were 33 miRILs described previously (Schmid et al. 2015). A file with the strains and the genetic map has been included (**Supplemental table S1 1**). Furthermore transgenic lines *amx-2(lf);Si[amx-2(Bristol)];let-60(gf)* and *amx-2(lf);Si[amx-2(Hawaii)];let-60(gf)* containing the N2- or CB4856-allele of *amx-2* in a *amx-2*(lf);*let-60*(gf) background were used in a confirmation experiment. For details regarding the construction of these lines, see (Schmid et al. 2015).

Strains were maintained on NGM agar seeded with OP50 bacteria at 20°C, strains were age-synchronized by bleaching and were harvested 72 hours post synchronization. One sample consisted of multiple pooled non-starved populations from ~4 separate 9 cm NGM plates.

### RNA isolation, cDNA synthesis, and cRNA synthesis

The RNA of the samples was isolated using the RNEasy Micro Kit from Qiagen (Hilden, Germany), following the ‘Purification of Total RNA from Animal and Human Tissues’ protocol with a modified lysing procedure. As prescribed, 150 µl RLT buffer and 295 µl RNAse-free water were used to lyse the samples, but with an addition of 800 µg/ml proteinase K and 1% ß-mercaptoethanol. This suspension was incubated for 30 minutes at 55°C and 1000 rpm in a Thermomixer (Eppendorf, Hamburg, Germany). Thereafter the protocol as supplied by the manufacturer was followed.

For gene-expression analysis 200 ng of RNA (as quantified by NanoDrop) was used in the ‘Two-Color Microarray-Based Gene Expression Analysis; Low Input Quick Amp Labeling’ -protocol, version 6.0 from Agilent (Agilent Technologies, Santa Clara, CA, USA).

### Microarray hybridisation and scanning

The Agilent *C. elegans* (V2) Gene Expression Microarray 4X44K slides were used to measure gene expression. As recommended by the manufacturer, the samples were hybridized for 17 hours and scanned by an Agilent High Resolution C Scanner. The intensities were extracted using Agilent Feature Extraction Software (version 10.7.1.1). The data was normalized using the Limma package in “R” (version 3.3.0 x64). For within-array normalization the ‘Loess’ method was used and the ‘Quantile’ method for between-array normalization (Zahurak et al. 2007; Smyth and Speed 2003). Unless mentioned otherwise, the log2 transformed intensities were used in subsequent analysis.

### Expanding the genetic map

Previously, the strains were genotyped with 73 fragment length polymorphisms (FLP) markers (as described in (Schmid et al. 2015; Zipperlen et al. 2005)) (Supplemental Table S2)

In order to increase the resolution of the genetic map, *cis*-eQTL of a large *C. elegans* eQTL experiment were used to further pinpoint the crossovers (Rockman et al. 2010). The gene expression of the miRILs was transformed to the mean of the two parental lines (R) by

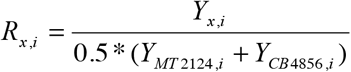

where Y stands for the untransformed intensities of strain x and spot i (1, 2, 3, …, 45220). The obtained values were correlated to the *cis*-eQTL effects from Rockman *et al*., 2010. Since the current study uses a newer version of the *C. elegans* microarray, only the spots that occurred in both designs were compared. The expression markers were generated following a procedure that is a variation of the method described in (West et al. 2006). Per 20 consecutive *cis*-eQTL, the correlation between the N2-eQTL effect and the transformed intensities (R) were calculated. In this way, the correlation would be negative if the genotype stemmed from CB4856 and positive if the genotype stemmed from N2.

In this way, per strain 424 gene expression markers were generated. In order to control for quality, the markers were filtered for calling the correct genotype in the four MT2124 replicas, the four CB4856 replicas, and in more than 50% of the samples an absolute correlation > 0.5 was required. This pruning resulted in 204 reliable gene expression markers. The quality of these markers was controlled by predicting the genotyped by expression of RIL AH2244, of which the gene expression was measured twice. There, we found that 10/204 selected gene expression markers were in disagreement of the genotype. This could be reduced to 0 by comparing only markers with an absolute correlation > 0.6. Therefore, all the correlations with an absolute value > 0.6 were assigned the predicted marker, which corresponds to an error rate < 0.01 per strain. The genotypes of the remaining markers were manually inferred from the surrounding markers. Furthermore, the genotypes at the ends of the chromosomes were inferred from the distal most assigned marker. This brings the size of the resulting map to 289 markers. The correlations and assigned genotypes can be found in **Supplemental table S3**.

### Evaluating the genetic map

The 289 marker set was analysed by correlation analysis for markers describing unique crossover events and to see if there are any strong linkages between chromosomes (**Supplemental figure S1**). This led to the selection of 247 markers indicating the border of a crossover event. To reduce the chances of false positives, we only used markers where the frequency of the least occurring genotype in the population was >15%. It has to be noted that this excluded most of chromosome IV (as the genotype was predominantly N2 due to selection for strains including the *let- 60(gf)* mutation (Schmid et al. 2015). Furthermore, also the *peel-1*/*zeel-1* region on chromosome I was excluded for the same reason (Seidel et al. 2008).

### eQTL mapping and threshold determination

Mapping of eQTL mapping was done in “R” (version 3.3.0 x64), the gene-expression data was fitted to the linear model,

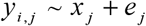

where y is the log2 normalized intensity as measured by microarray of spot i (i = 1, 2, …, 45220) of miRIL j. This is explained over the genotype (either CB4856 or N2) on marker location x (x = 1, 2, …, 247) of miRIL j.

A permutation approach was used to determine an empirical false discovery rate (Snoek et al. 2017; Vinuela et al. 2010). First, the log2 normalized intensities were randomly distributed per gene over the genotypes. This randomized dataset was tested using the eQTL mapping model. This procedure was repeated 10 times. The threshold was determined using

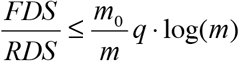

Where, at a specific significance level, the false discoveries (FDS) were the averaged outcome of the permutations and read discoveries (RDS) were the outcome of the eQTL mapping. The value of m_0,_ the number of true null hypotheses tested, was 45220-RDS and for the value of m, the number of hypotheses tested, the number of spots (45220) was taken. Because this study only used a limited set of strains, a more lenient threshold of 0.1 was taken as the *q*-value (Benjamini and Yekutieli 2001), which resulted in a threshold of –log10(p) > 3.2.

### Statistical power calculations

The statistical power of the mapping was determined at the significance threshold of –log10(p) > 3.2. Using the genetic map (33 strains with 247 markers), 10 QTL were simulated per marker location, explaining 20-80% of the variation (in increments of 5%). Next to a peak, also random variation was introduced, based on a standard normal distribution (mu = 0 and sigma = 1). Peaks with corresponding explanatory power were simulated in this random variation (e.g. a peak size of 1 corresponds to 20% explained variation). Furthermore, also random data without peaks was generated. This simulated data was mapped as described above and for each simulated peak it was determined: (i) if it was detected correctly, (ii) how precise the effect size estimation was, and (iii) how precise the location was determined. An overview of the results can be found in **Supplemental table S4**.

### eQTL analysis

The distinction between *cis-* and *trans-*eQTL was made on the distance between the gene and the eQTL-peak. Given the relatively low number of unique recombinations (due to the limited set of strains), the *cis*-eQTL window was set at 2 Mb. This means that if a gene lies within 2 Mb of the QTL peak or the confidence interval, it is called a *cis-*eQTL. The confidence interval was based on a –log10(p) drop of 1.5 compared to the peak.

*Trans-*bands were identified based on a Poisson-distribution of the mapped *trans*-eQTL (as in (Rockman et al. 2010)). The number of *trans*-eQTL was counted per 1 Mb bin, leading to the identification of 52 bins with *trans*-eQTL. Since we mapped 1149 *trans*-eQTL peaks (spots) to 52 bins, we expected 22.10 *trans*-eQTL per bin with *trans*-eQTL assigned to it. Based on a Poisson-distribution, any bin linking more than 30 spots with a *trans*-eQTL would have a significance p < 0.05 and more than 38 *trans*-eQTL would yield a significance of p < 0.001. In this way, 6 *trans*-band loci were identified.

Comparative analysis against previous eQTL experiments was done on re-mapped experiments downloaded from WormQTL (Snoek et al. 2013; van der Velde et al. 2014) and EleQTL (http://www.bioinformatics.nl/EleQTL). This dataset consists of 4 different experiments representing 9 different conditions. The first set contains eQTL in two temperature conditions, 16°C and 24°C, measured in the L3 stage (Li et al. 2006). The second set contains eQTL over 3 life stages: L4 juvenile animals grown at 24°C, reproducing adult animals (96h) grown at 24°C, and aging animals (214h) grown at 24°C (Vinuela et al. 2010). The third set contains eQTL from a single experimental condition (young adults grown at 20°C) measured on a large RIL panel (Rockman et al. 2010). The fourth set contains eQTL from three experimental conditions over the course of a heat-shock treatment: a control condition (L4 animals grown for 48h at 20°C), a heat-shock condition (L4 animals grown for 46h at 20°C and exposed to 2h of 35°C), and a recovery condition (similar to heat-shock, only followed by 2h at 20°C) (Snoek et al. 2017). Each of these experiments was compared at FDR = 0.05 to the eQTL mapped at FDR = 0.10 in the *let-60(gf)* sensitized miRILs.

Since the dataset of Rockman *et al*., 2010 was used for expanding the genetic map, analysis was also conducted excluding this study. The reason for excluding this set from analysis was that a bias could be introduced in overlap with the *cis*-eQTL. The main conclusion that *cis*-eQTL are less unique than *trans*-eQTL also stands with this analysis. Excluding the Rockman *et al*. data, resulted in detection of 999/1516 (65.9%) *cis*-eQTL and 171/898 (19.0%) *trans*-eQTL present in previous experiments.

### Enrichment analysis

Gene group enrichment analysis was done using a hypergeometric test on the unique genes (not on the spots) with the following criteria: Bonferroni corrected p-value < 0.05, size of the category n>3, size of the overlap n>2.

The following databases were used: The Wormbase (www.wormbase.org) WS220 gene class annotations, the WS256 GO-annotation, anatomy terms, phenotypes, RNAi phenotypes, developmental stage expression, and disease related genes (Harris et al. 2014); the MODENCODE release 32 transcription factor binding sites (www.modencode.org; (Gerstein et al. 2010; Niu et al. 2011)), which were mapped to transcription start sites (according to (Tepper et al. 2013)); the KEGG pathway release 65.0 (Kyoto Encyclopedia of Genes and Genomes (www.genome.jp/kegg/) (Ogata et al. 1999).

### amx-2 allelic comparison

Four independent transgenic, 2 of *amx-2(lf);Si[amx-2(Bristol)];let-60(gf)* and 2 of *amx-2(lf);Si[amx-2(Hawaii)];let-60(gf)*, were used to investigate the effect of the *amx-2* CB4856 and N2 allele on gene expression. These lines contain single-copy insertions on LGII of the N2- or CB4856-alleles of *amx-2* in an N2 *amx-2*(lf); *let-60*(gf) background. The effect of different alleles on gene expression was measured by micro-array for each independent transgenic and compared to the effects of the eQTL mapping to the *trans*-band closest to the position of *amx-2*.

### Connectivity network analysis

To investigate the connectivity and function of the affected genes we used WormNet (version 3) (Cho et al. 2014) and GeneMania pluging for Cytoscape (version 3.4.0 (Montojo et al. 2010; Shannon et al. 2003)). WormNet was used to investigate enrichment in connectivity within groups of genes. For example, groups of co-expressed genes mapping to the same *trans*-band were assumed to share the same regulator. Further evidence for this co-expression and regulation can be found in WormNet if these genes are significantly more connected than by chance. GeneMania was used to find the closest neighbours, those genes with the most connections, of *amx-2* and *let-60* to locate the genes by which *amx-2* modifies RAS signalling.

## RESULTS

### Expression-QTL mapping in miRIL population

Using microarrays, we measured gene expression levels in 33 miRILs sensitized for RAS/MAPK signalling by introgression of a *let-60* (gf) mutation in a segregated N2/CB4856 genetic background (**Supplemental figure S2**) (Schmid et al. 2015). Analysis of the statistical power of this population with a genetic map of 247 informative markers showed that we can detect 77% of the QTLs explaining 40% of the variation (**Supplemental table S4**). We detected 2303 genes (represented by 3226 array spots) with at least one eQTL (FDR=0.1, -log10(p)>3.2; **Table 1, Figure 1, Supplemental table S5**). For 1516 of these genes a *cis-*eQTL was found, indicating regulation from within a window of 2 Mb around the affected gene. For 898 genes a *trans-*eQTL was found, showing regulation more distal from the affected transcript. Most *cis-*eQTL (1074 out of 1516; ~71%) show a positive effect for the N2 allele, whereas for the *trans-*eQTL the positive and negative effects were equally distributed over the N2- and CB4856-allele (**Table 1**).

**Table 1.** The number of genes with an eQTL, in brackets the number of spots.

|  | *cis*-eQTL | *trans*-eQTL |
| --- | --- | --- |
| N2 higher | 1074 (1564) | 448 (582) |
| CB4856 higher | 456 (632) | 464 (576) |
| Total<sup>1</sup> | 1516 (2196) | 898 (1149) |

<sup>1</sup>: The discrepancy between the sum of N2 higher and CB4856 higher and the total is due to genes being represented by multiple spots, which often represent different splice variants.

**Figure 1.**
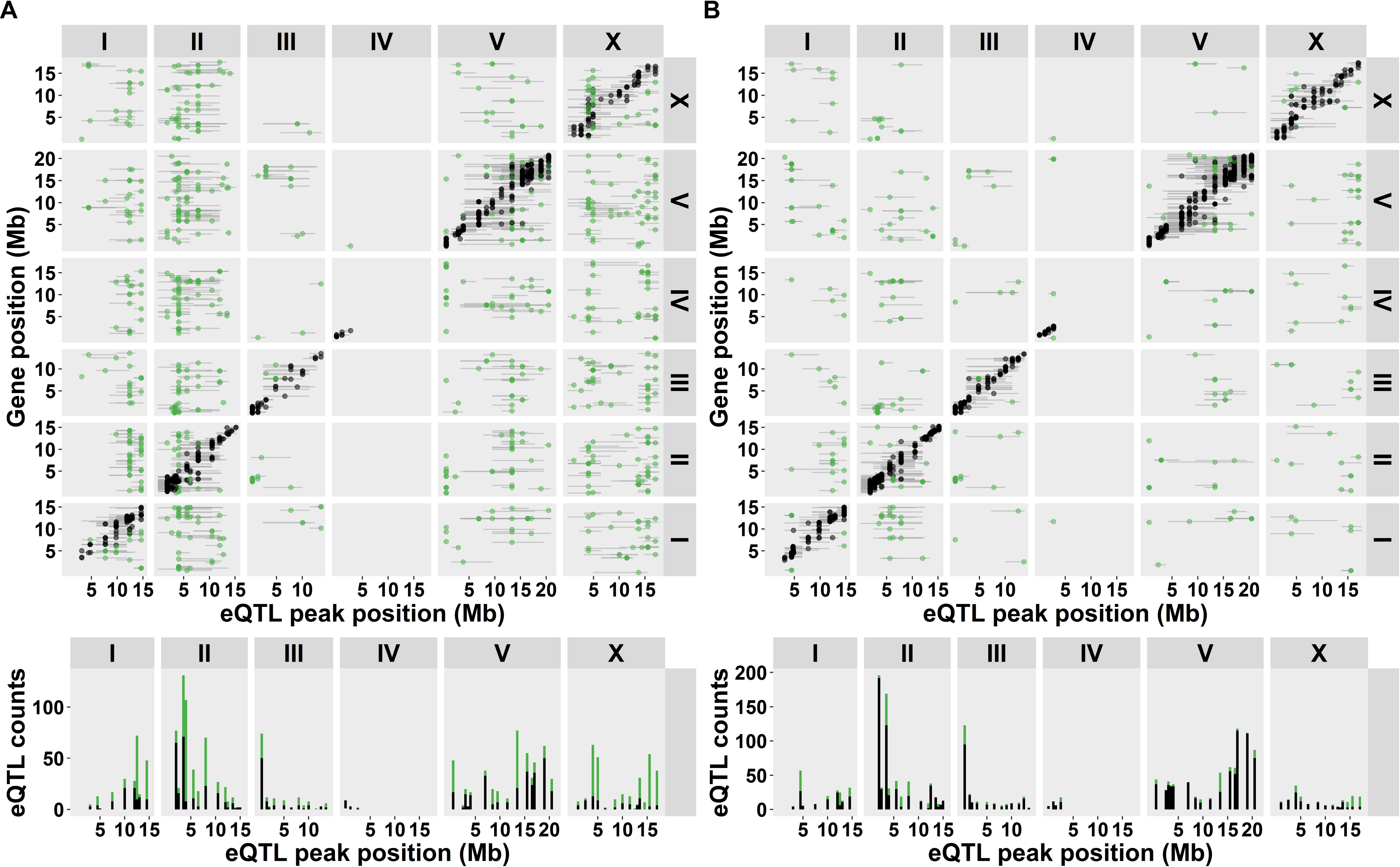
eQTL mapping in the *let-60(gf)* miRIL population. On top the *cis-trans* plot is shown. On the horizontal axes the positions of the eQTL are plotted, along the six chromosomes. The location of the genes with an eQTL is plotted on the vertical axis. The black dots represent *cis*-eQTL (lying within 2 Mb of the regulated gene), the green dots represent *trans*-eQTL (eQTL lying elsewhere). The grey horizontal lines indicate the confidence interval of the QTL location (based on a 1.5 drop in – log10(p)). The bottom histogram shows the number of eQTL per genomic location. Note that on chromosome IV hardly any eQTL are mapped, this is because of linkage of the *let-60(gf)* mutation (which is located at IV:11.7 Mb).

The eQTL were not equally distributed across the genome. We detected different clustered *trans*-eQTL as “hotspots” or *trans*-bands. *Trans*-bands are frequently found in eQTL experiments and indicate a locus with one or multiple alleles affecting the expression of many distant genes. To identify the *trans*-bands in our experiment, we calculated the chance of overrepresentation of *trans*-eQTL per marker, using a Poisson distribution (as in (Rockman et al. 2010)). This method was applied per 1 Mb window on the genome. The *trans*-bands that were identified in adjacent windows were merged. This way, 6 *trans*-bands were identified (Poisson distribution, p < 0.001; **Table 2**).

**Table 2.**
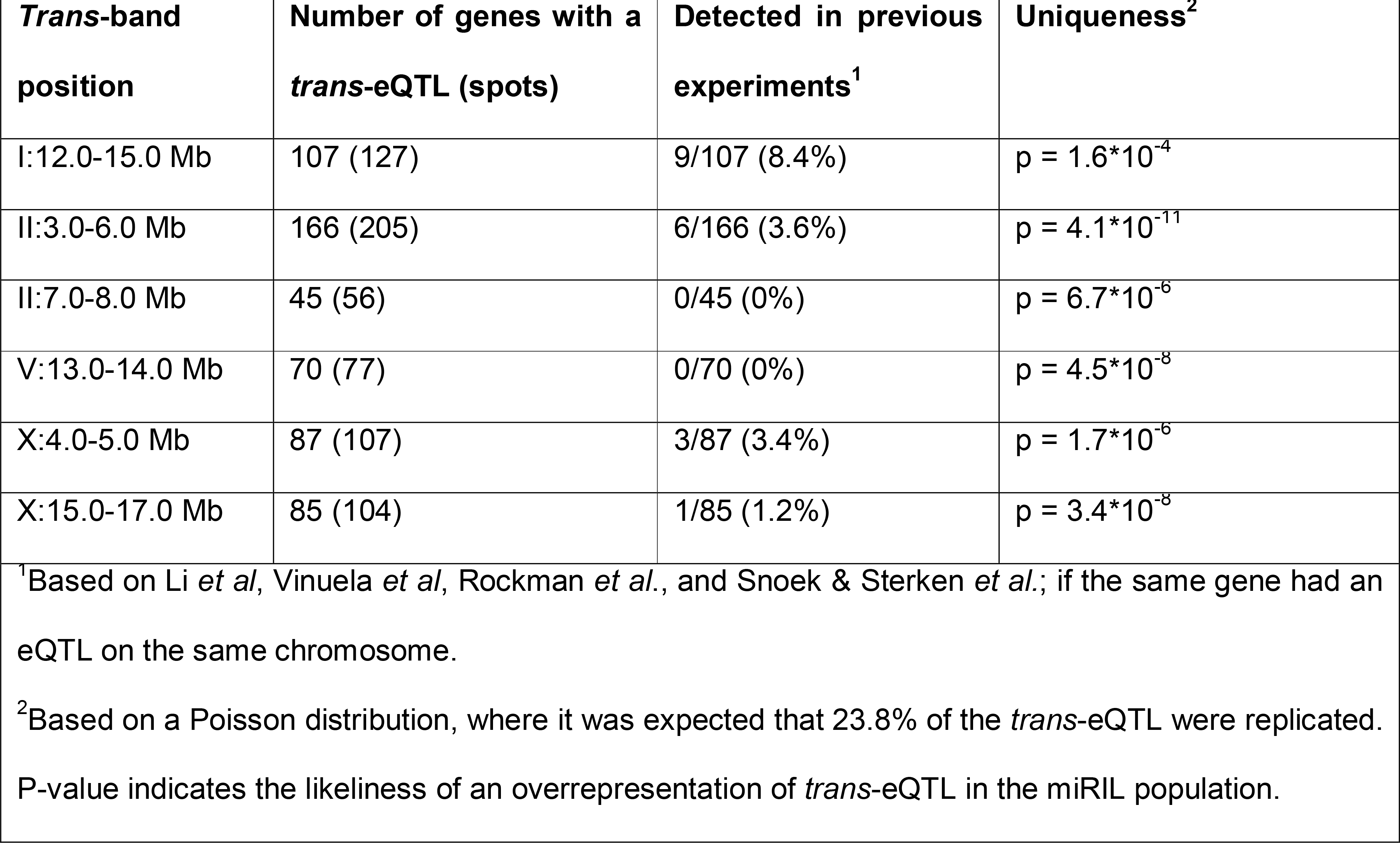
trans-bands detected in the sensitized miRILs.

### Specificity of let-60 eQTL

By comparative analysis of different environments and/or populations, it has been shown that eQTL *trans*-bands can be population, environment, or age specific (for example, see (Li et al. 2006)). To investigate if and which eQTL are specifically found in the sensitized miRILs, we compared the detected eQTL to the eQTL found in non-sensitized RILs (Li et al. 2006; Vinuela et al. 2010; Rockman et al. 2010; Snoek et al. 2017). These datasets contain eQTL mapped over 9 different conditions (see Methods), which were compared at an FDR = 0.05 with the eQTL mapped in the sensitized miRILs. We found that ~73% (1112 out of 1516) of the genes with a *cis*- eQTL in the miRIL population were also found in one or more of the previous studies (**Figure 2, Supplemental figure S3B**). When taking the allelic effects in account, of the 1074 genes with a *cis-*eQTL that gave a higher expression linked to the N2 allele, 816 (~76%) were found back in other studies. For the 456 genes with a *cis-*eQTL that gave a higher expression linked to the CB4856 allele, 310 (~68%) were found back in other studies. This shows that the majority of the *cis*-eQTL detected in the miRIL population can be also be detected in other experiments.

**Figure 2.**
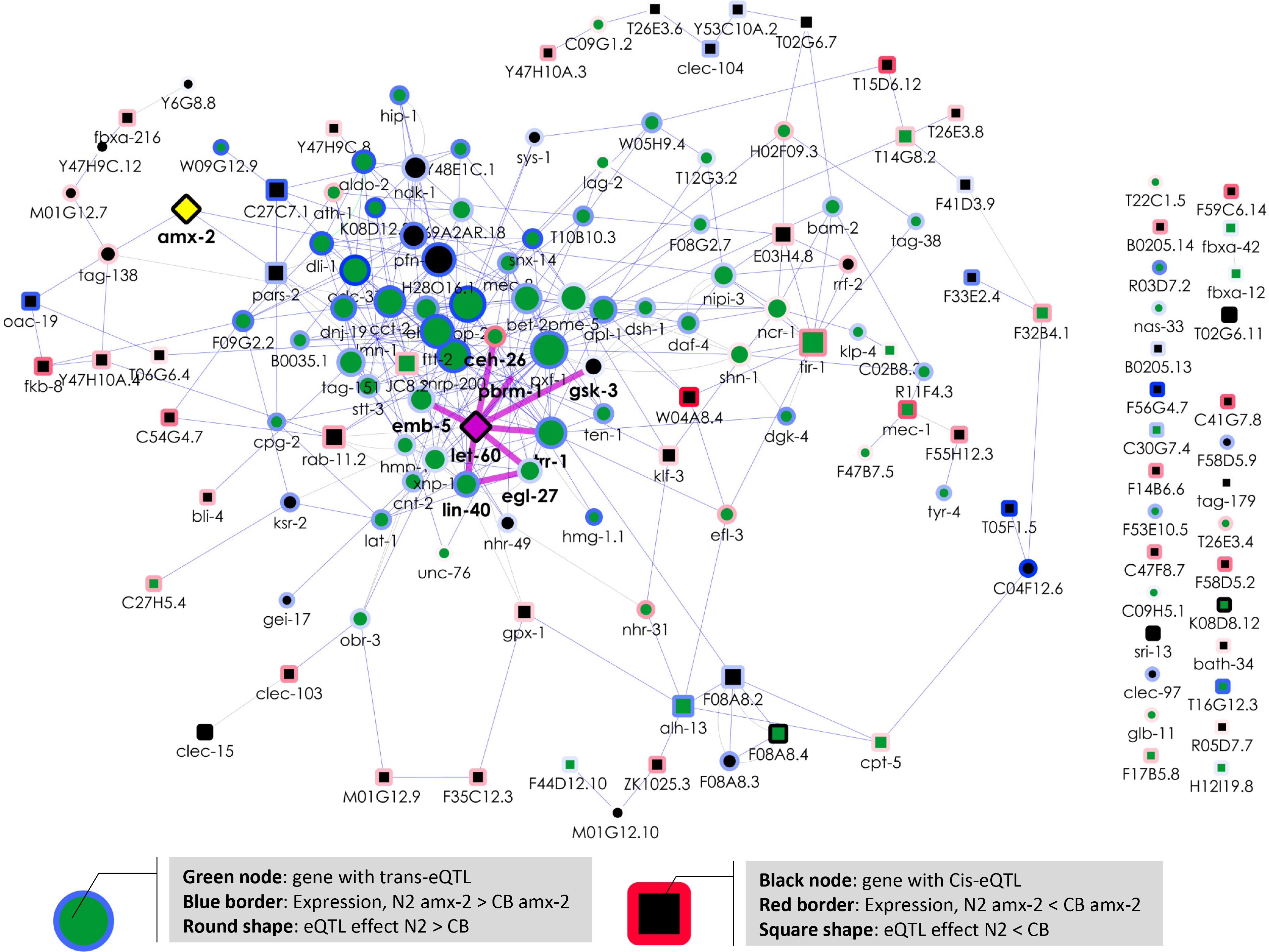
eQTL detected in previous experiments. The eQTL mapped in the *let- 60(gf)* RIL population were compared to eQTL mapped in three published *C. elegans* eQTL studies, representing 9 different conditions (Li et al. 2006; Vinuela et al. 2010; Rockman et al. 2010; Snoek et al. 2017). The histogram shows how many of the eQTL mapped in this study were found back in these independent conditions. Whereas the majority of *cis*-eQTL was mapped in previous studies, only a minority of the *trans*-eQTL was.

In contrast, for the *trans-*eQTL detected in our experiment - not taking location into account - we only found ~24% (214 out of 898) of the genes had a *trans-*eQTL in at least one of the previous experiments (**Figure 2** and **Supplemental figure S3C**). In order to further compare the *trans*-eQTL overlap, it was investigated whether the *trans*-eQTL co-localized on the same chromosome. Using that criterion, ~9% (80 out of 898) of the genes with a *trans*-eQTL were found in a previous study. This showed that the majority of the genes with a *trans*-eQTL were specifically detected in the sensitized miRIL population. This observation also extended to the *trans*-bands. To test the specificity of the *trans*-bands, we counted the number of times a gene within the miRIL *trans*-bands had a co-locating *trans*-eQTL in other studies. All *trans*-bands were found to be specific for the *let-60(gf)* miRILs (Poisson distribution, p < 0.001; **table 2**). This provides additional evidence that these *trans*-bands are specific for interaction between the genetic background and the *let-60(gf)* mutation.

Together, we identified 404 genes with a novel *cis*-eQTL and 684 genes with a novel *trans*-eQTL (**Figure 3A**). As our method can be biased by the way the genetic map was constructed (using expression differences, see **Materials and methods**), we conducted the same analyses using the original FLP map. Our conclusions on the detection of novel *cis*- and *trans*-eQTL were not affected depending on the genetic map (for details, see **Supplemental text S1**). This implies that a substantial majority of *trans*-eQTL showed a mutation dependent induction. This is also illustrated by showing the eQTL that were consistently found (**Figure 3B**), which mainly shows the *cis*-diagonal and a few *trans*-eQTL.

**Figure 3.**
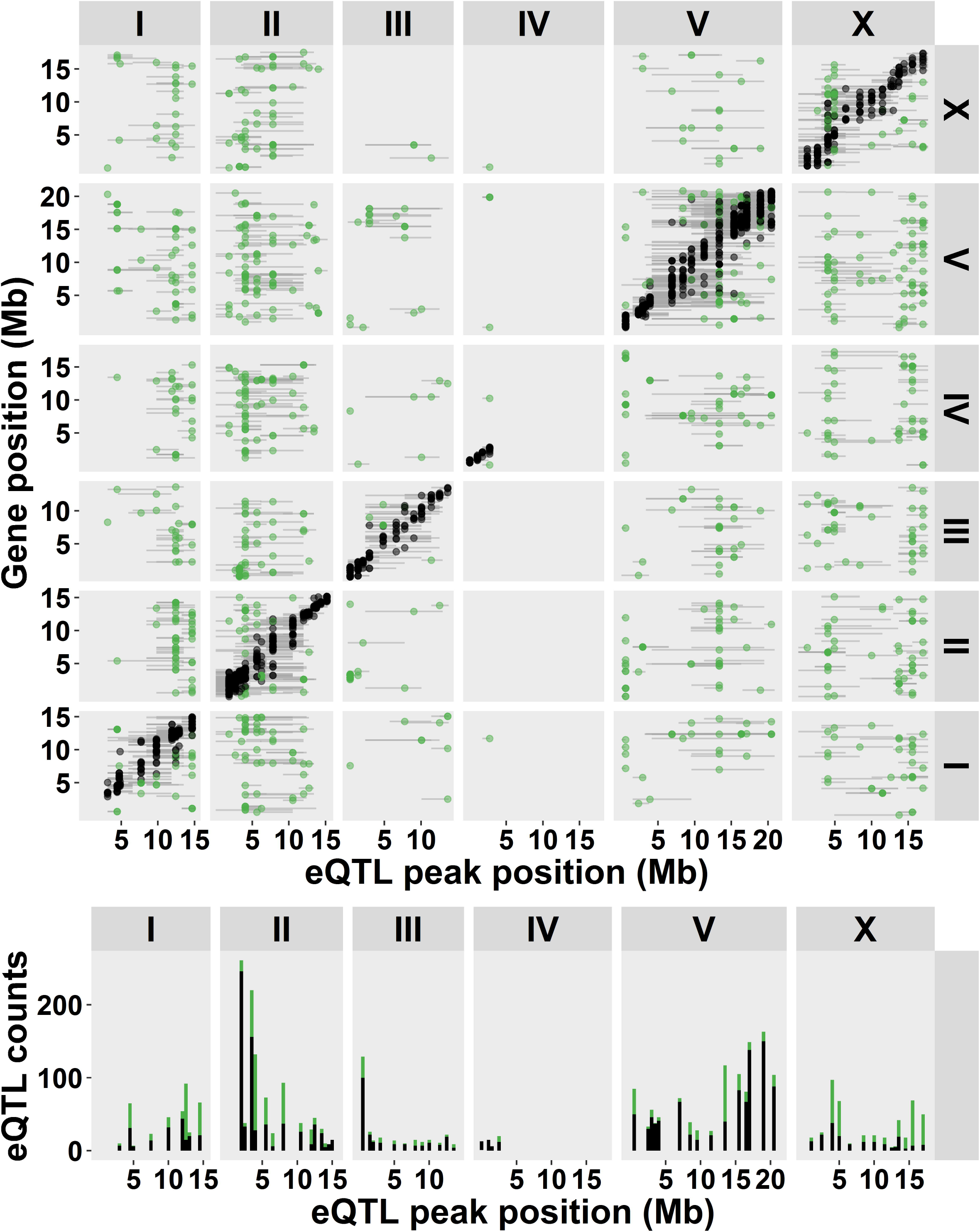
eQTL mapped in the *let-60(gf)* miRIL population compared to other studies (Li et al. 2006; Vinuela et al. 2010; Rockman et al. 2010; Snoek et al. 2017). As in **figure 2**, the *cis*- and *trans*-eQTL are plotted. (**A**) eQTL only found in the miRIL population. (**B**) eQTL found over multiple studies.

### Functional allelic differences

Enrichment analysis on groups of genes with an eQTL mapping to the same *trans*- band can be used to uncover the functional consequences of the allelic variation at the selected locus. Enrichment analysis was done on GO-terms, KEGG, Anatomy-terms, disease phenotypes, development expression, phenotypes, transcription-factor binding sites, and RNAi phenotypes (**Supplemental table S6**).

The genes with *cis*-eQTL were enriched for genes in the gene classes: *math*, *bath*, *btb*, and *fbx*. These classes contain many genes that function as intra- and extra-cellular sensors involved in gene-environment interactions and were found highly polymorphic between N2 and CB4856, but also between other wild-isolates (Thompson et al. 2015; Volkers et al. 2013). The genes with *trans*-eQTL with higher expression linked to CB4856 loci were enriched for genes related to development in general and gonad development specifically. These enrichments were likely to stem form the *trans*-bands on chromosome I:12-15 Mb and II: 3-6 Mb specifically. These two *trans*-bands mainly consisted of genes with a higher expression linked to the CB4856 locus and were enriched for genes associated with reproduction and/or development.

### Allelic effect of amx-2 on gene expression

The *trans*-band on chromosome I:12-15Mb co-localizes with a previously identified QTL for vulval induction, for which *amx-2* was identified as polymorphic regulator (Schmid et al. 2015). To determine if the *trans*-eQTL mapping to the *amx-2* locus were affected by the allelic difference between N2 and CB4856, we measured gene expression in transgenic lines containing single-copy insertions of either allele of *amx-2* in a *amx-2(lf)*; *let-60(gf)* background. For each allele, two independent strains were created and the transcriptome was measured using microarrays (Schmid et al. 2015). From these measurements the allelic effects of *amx*-2 on gene expression were determined and compared with the *trans*-eQTL effects measured in the miRIL population. These were found to correlate significantly (**Supplemental figure S4**; R^2^ ~0.34; p < 2*10^-8^). Specifically, 61 of the 77 genes with an eQTL co-localizing with *amx-2* showed the same allelic effect in the transgenic strains carrying the two *amx-2* variants. Furthermore, analysis of the effect directions showed that the *trans*-band on chromosome I (**Table 2**) consists of two separate *trans*-bands (see **Supplemental figure S4**); the first co-localizing with *amx-2* and the second lying more distal on chromosome I.

To further investigate the mechanism by which *amx-2* modifies *let-60* Ras/MAPK signalling, we used the well-established *C.elegans* connectivity networks WormNet (Cho et al. 2014) and GeneMANIA (Montojo et al. 2010; Shannon et al. 2003). WormNet showed that the genes mapping to the *amx-2* locus were more connected than expected by chance (p < 3*10^-13^). Visualising the connectivity of these genes together with *amx-2* and *let-60* using GeneMANIA identified one major gene-cluster (**Figure 4**). As expected, genes affected by the *trans*-band were found to be highly connected and at the core of the cluster, indicating a shared biological function. In contrast, 25% of the genes with a *cis*-eQTL at the *trans*-band location were not connected and those that were connected had only one to three connections. Interestingly, *let-60* was at the core of the cluster of genes with a *trans*- eQTL, whereas *amx-2* was at the periphery. We found six genes, *lin-40* (also called *egr-1*), *egl-27*, *trr-1*, *pbrm-1*, *ceh-26* (also called *pros-1*) and *emb-5*, previously found to have a genetic interactions with *let-60* (Lee et al. 2010; Lehner et al. 2006; Byrne et al. 2007). These could be the genes through which *amx-2* affects RAS/MAPK signalling.

**Figure 4.**
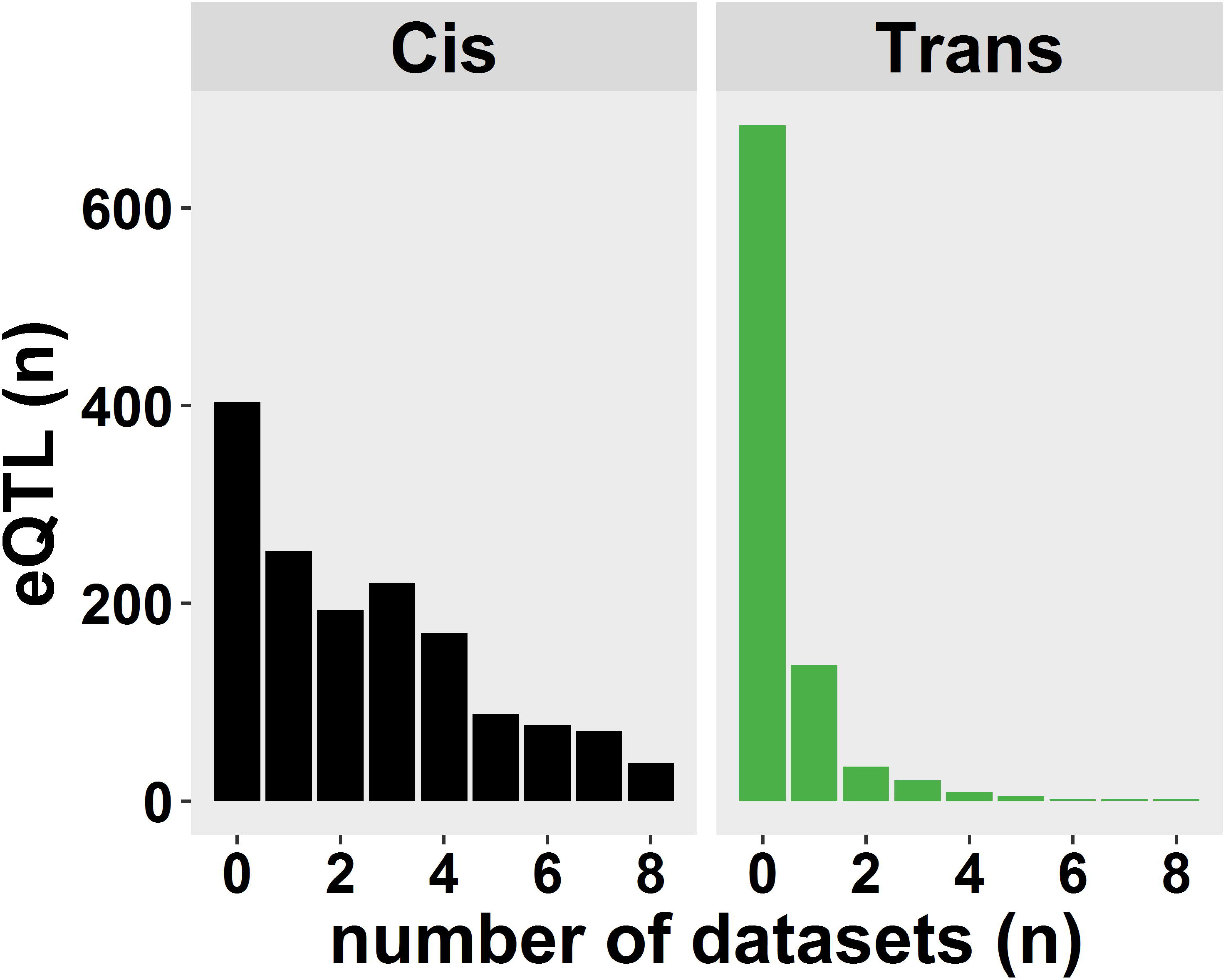
Interaction network of genes co-localizing with *amx-2*. Genes with an eQTL co-localizing with *amx-2* were selected together with *amx-2* (yellow) and *let-60* (pink). Interactions were obtained from Genemania (Montojo et al. 2010; Shannon et al. 2003). Node colour indicates: *cis* in black and *trans* in green. Node shape indicates eQTL effect, N2 > CB4856 as a circle and CB4856 > N2 as a square. Node border colour gradient indicates expression in *amx-2* transgenic lines (blue N2-*amx-2*-allele > CB4856-*amx-2* allele, red CB4856-*amx-2* allele > N2 *amx-2* allele). Node size indicates number of edges connected. Edges in blue show co-expression links and edges in pink show genetic interactions as found by (Lee et al. 2010; Lehner et al. 2006; Byrne et al. 2007). Genes connected by genetic interactions have bold gene names (as well as *amx-2*).

## DISCUSSION

Ras/MAPK signalling is strongly affected by gain of function mutations in *let-60* as shown by the strong effect on vulva development in *C. elegans* (Beitel et al. 1990). The allele *let-60(n1046)* hyperactivates Ras/MAPK signalling, leading to a multivulva phenotype and the differentiation of more than three vulval precursor cells. In the miRIL population, which carries a *let-60(n1046)* mutation in a segregating N2/CB4856 background, a wide range of vulval induction (VI) was found. While in the full N2 background VI of the *let-60(gf)* mutation was 3.7 induced vulval cells on average, the VI in the miRIL population varied from 3.0 to 5.7 induced vulval cells, illustrating the strong modulatory effect of the genetic background on the mutant phenotype (Schmid et al. 2015). By measuring gene expression across the different miRILs, we gained insight into the genetic background effects on the transcriptional architecture of *let-60(gf)* sensitized miRILs.

### The let-60(gf) mutation affects gene expression in trans

To our knowledge this is the first study where a mutation, introgressed into a panel of RILs, is used to investigate the genetic architecture of transcript variation by eQTL analysis. Unfortunately, the direct effect of the *let-60(gf)* allele on the genetic background is impossible to determine in this population, since there is no population with the same genetic background lacking the *let-60(gf)* mutation. Therefore we used previously published eQTL studies in *C. elegans* as estimation for the miRIL specific eQTL. We found that introgression of *let-60(gf)* leads to an overrepresentation of specific *trans*-eQTL, while the *cis*-eQTL showed a high overlap with eQTL observed in the absence of *let-60(gf)* allele. Although our study is based on a relatively small number of strains, the verification of identified eQTL against other eQTL studies in *C. elegans* shows that at least 73% of the identified *cis-*eQTL were discovered before. This supports our power analysis, showing that our study has enough statistical power to detect eQTL with larger effects.

Given that *cis*-eQTL are most likely genes with polymorphisms in or near the gene itself, it is expected that most are discovered before. In support, most of the *cis*- eQTL genes show higher expression in N2, making it likely that a relatively large part of these *cis*-eQTL indeed stem from polymorphisms causing mis-hybridization on the microarrays instead of true expression differences (Rockman et al. 2010; Alberts et al. 2007). Therefore, most of the *cis*-eQTL will not represent gene expression changes that can be linked to the *let-60(gf)* mutation in the miRIL population. That is also exactly what we found when constructing the regulatory network by connecting *cis*- and *trans*-eQTL to *let-60*; *cis*-eQTL are only loosely - or not at all - connected to the network.

Compared to *cis*-eQTL, the *trans*-eQTL identified in the miRIL population were hardly found in previous eQTL studies in *C. elegans* (only 24% was detected previously). However, it should be noted that *trans*-eQTL on themselves are also environment- and age-dependent (Vinuela et al. 2010; Francesconi and Lehner 2014; Li et al. 2006; Snoek et al. 2017), therefore it possible that not all the *trans*- eQTL are indeed *let-60* induced. For example, they might be age-related (Vinuela et al. 2010). Another aspect that influences detection of *trans*-eQTL is the power of the study, as *trans*-eQTL explain less of the variation than *cis*-eQTL. However, by being stringent in the overlap (requiring only one match over nine different environments) it is likely that most of these eQTL would be detected. This makes it likely that many of the identified *trans*-eQTL are specifically linked to an effect of the *let-60(gf)* mutation in the genetic background. This extends the idea that environmental perturbations can reveal additional genetic variation (Li et al. 2008), by perturbing the genetic environment (Kammenga et al. 2008; Duveau and Felix 2012; Schmid et al. 2015). The advantage of mutational perturbation and perturbation via induced responses in eQTL studies (for example, see (Snoek et al. 2012) and (Snoek et al. 2017)), is the placement of the transcriptional response into a context. Yet, as the phenotypic influence of *let-60(gf)* in the miRIL population was investigated before, we know it results in a phenotype In the study presented here, the context is the *let-60(gf)* mutation, which results in a phenotype where vulval induction is affected. It is therefore interesting that the *trans*-bands co-localize with QTL mapped for index VI index in the same miRIL population: *trans*-band I:12-15 Mb overlaps with QTL1b, *trans*-band II:7-8 Mb with QTL2, and *trans*-band V:13-14 Mb with QTL3 (Schmid et al. 2015). This co-localization adds to the plausibility that the novel *trans*-eQTL detected in the miRIL population are indeed due to the *let-60(gf)* mutation and its interaction with the genetic background.

### A trans-band on chromosome I links to variation in the amx-2 gene

Using the same *let-60(gf)* miRIL population, we have previously identified the polymorphic monoamine oxidase *amx-2* as a background modifier that negatively regulates the Ras/MAPK pathway and the VI phenotype (Schmid et al. 2015). The *trans*-band on chromosome I co-localizes with the *amx-2* QTL for VI (QTL1b, (Schmid et al. 2015)), it is therefore possible that the modifier *amx-2* also affects gene expression. More specifically, the *trans*-band on chromosome I originally identified in the miRILs consists of two separate regulatory loci. The N2 allelic effect at the *amx-2* locus is mostly negative, whereas for the more distal part of the original *trans*-band the N2 allelic effect is mostly positive. Yet, it seems unlikely that *amx-2* is a direct gene expression effector. As *amx-2* is a mitochondrial monoamine oxidase type A, a catalyser of neuropeptide oxidative deamination, it will probably not influence gene expression directly (Tipton et al. 2004). Placing the *trans*-eQTL of the *amx-2 trans-*band in a gene interaction network supports this line of reasoning: although *let-60* is in the centre of the *trans*-eQTL, *amx-2* is only peripheral.

By measuring gene expression in transgenic lines expressing the N2 or CB4856 allelic variants of *amx-2* in an *amx-2(lf);let-60(gf)* genetic background, we attempted to link the allelic effect of *amx-2* to the *trans*-eQTL mapping to the *amx-2* locus. As a significant correlation between the expression differences in the transgenic lines and the *trans*-eQTL effects mapping to the *amx-2* locus is found, these *trans*-eQTL can be confidentially linked to allelic variation in *amx-2*. As discussed before, it is unlikely that *amx-2* is the direct cause of the transcriptional variation, but rather acts through an indirect mechanism. One route of effect might be through *amx-2* mediated degradation of serotonin (5-HT) to 5-HIAA, which both affect VI (as discussed in (Schmid et al. 2015)). Subsequently, the affected VI might result in different gene expression levels. It is interesting to remark that the *amx-2 trans*- band is characterized by down-regulation of expression related to the N2 genotype as well as a decreased VI for that genotype.

As this places *amx-2* in the causal chain of events, we think it is most likely that *amx-2* does not affect gene expression directly but via another gene involved in Ras/MAPK signalling, possibly directly linked to *let-60*. Therefore, we hypothesize that *amx-2* affects Ras/MAPK signalling via one or more of the six previously found genes with a genetic interaction with *let-60*: *lin-40* (also called *egr-1*), *egl-27*, *trr-1*, *pbrm-1*, *ceh-26* (also called *pros-1*), and *emb-5*. For example, *egl-27* and *lin-40* both have a *trans*-eQTL mapping to the *amx-2* locus, and are the two MTA (metastasis-associated protein) homologs found in *C. elegans* (Herman et al. 1999; Ch'ng and Kenyon 1999; Solari et al. 1999). The proteins act in the NURD chromatin remodelling complex, which has previously been shown to antagonize Ras-induced vulval development (Solari and Ahringer 2000). Chromatin remodelling provides a more likely mechanism of action than the molecular role of *amx-2* itself.

### Implications for understanding the Ras/MAPK pathway

How can our results help a better understanding of the Ras/MAPK pathway? Our previous investigation on VI in the miRIL population resulted in the identification of three QTL harbouring polymorphic Ras-signalling modifiers. Expanding our research to genetic variation affecting gene expression in a *let-60(gf)* sensitized RIL population uncovered six *trans*-bands (or ‘eQTL hotspots’). As mentioned before, these *trans*- bands overlap with the QTL mapped for VI in Schmid *et al*., 2015. There are three more *trans*-bands that do not overlap with QTL for VI, but are also likely to represent modifier loci of the Ras/MAPK pathway, possibly underlying other Ras/MARK associated phenotypic differences.

The main advantage of studying the genetic architecture of gene expression in the miRIL population is that it creates more insight in the genes and pathways affected by allelic variation acting on the Ras/MAPK pathway. It identifies hidden genetic variation; genetic variants that are unlocked under altered environmental conditions or when the genetic background is modified (for a review, see (Paaby and Rockman 2014) and (Kammenga 2017)). Identification of background modifiers of disease pathways is important for gaining insight into individual based differences of disease contraction. Mutant phenotypes can be strongly affected by the genetic background. For example, large variation in traits between different backgrounds in many different mutated genes has been observed in *C. elegans* (Vu et al. 2015; Paaby et al. 2015; Duveau and Felix 2012). A major discovery was that this variation in mutant phenotypes could be predicted from gene expression variation (Vu et al. 2015). Those results are in line with the discovery of *trans*-bands at the location of each VI QTL. Furthermore, results presented here (and previously (Elvin et al. 2011)) link variations in individual gene expression levels to enhancing or diminishing the severity of a Mendelian disorder caused by a so-called “major gene”. More specifically, these differences seem to stem from a couple of modifier loci harbouring polymorphic regulators.

So far, we only explored the *amx-2 trans*-band, and it is likely that the other loci also contain polymorphic modifiers of the Ras/MAPK pathway. Genetic perturbation by *let-60(gf)* leads to a strong increase in specific *trans*-acting eQTL organized over six *trans*-bands, thus supporting the involvement of multiple genes as modifiers. Given the fact that we detected *amx-2* both via an indirect transcriptional response, but also mechanistically, we feel confident that the detected loci indeed harbour other modifiers. A single gene mutation apparently has the capacity to unlock a huge number of novel interactions controlled by many genes across different genetic backgrounds, further studies should be conducted to identify and characterize these underlying polymorphic regulators.

## CONCLUSION

Introgression of a mutation in a segregating genetic background allows for identification of polymorphic modifier loci. By introducing a *let-60(gf)* mutation in a N2/CB4856 genetic background, in the form of the miRIL population and measuring the transcriptome, we identified six *trans*-bands specific for the *let-60(gf)* miRIL population. The majority of *trans*-eQTL are specific for the miRIL population, showing that genetic variation in gene expression can be specifically modified by a background mutation. Therefore, genetic perturbation can be viewed as analogous to environmental perturbation, which also results in specific *trans*-eQTL. We demonstrated the involvement of *amx-2* and the allelic variation between N2 and CB4856 in gene expression variation originating from chromosome I. Yet, we think it is unlikely that *amx-2* directly affects gene expression variation. Instead, we prefer the hypothesis that allelic variation in *amx-2* indirectly affects gene expression, possibly through the NURD complex.

## AVAILABILITY OF DATA AND MATERIALS

All strains used can be requested from the authors. The transcriptome datasets generated and the mapped eQTL profiles can be interactively accessed via http://www.bioinformatics.nl/EleQTL. Moreover, the raw transcriptomic data is also available at ArrayExpress (E-MTAB-5856)

## SUPPLEMENTAL DATA

**Supplemental text S1**: Text explaining the analysis of eQTL in the original FLP-based genetic map.

**Supplemental figure S1:** Between-marker correlations of the genotypes in the *let- 60(gf)* sensitized RIL population. Per marker pair, the correlations are shown in a heat-plot. On the x-axis marker 1 and on the y-axis marker 2 is shown. Most in-between chromosome correlations are |R|< 0.6. However, between chromosome I:2.6-2.7 Mb and chromosome IV3.6-4.2 Mb and chromosome II:3.1-3.4 Mb and chromosome III:0.0-1.4 Mb higher correlations were detected.

**Supplemental figure S2:** A figure of the genetic map of the sensitized RIL population. On the x-axis the physical position on the genome is given an don the y-axis the genotypes used in this study. The genotypic origin at each genomic position is indicated by colour (orange for N2 and blue for CB4856).

**Supplemental figure S3:** Comparison of eQTL mapped in the *let-60(gf)* miRIL population with previous genetical genomics studies in *C. elegans*. (**A**) eQTL effect sizes per condition per study, split out for *cis*- and *trans*-eQTL. (**B**) Constitutively found *cis*-eQTL. On the x-axis the QTL position in the sensitized RIL population is shown, on the y-axis the QTL position in the compared experiment is shown. Therefore, a point on the diagonal indicates a QTL mapped to the same position. The lines indicate the 1.5 LOD-drop confidence interval of the QTL position. (**C**) Constitutively found *trans*-eQTL, annotations are the same as in (**B**).

**Supplemental figure S4:** Comparison between eQTL effects on chromosome I and the difference between N2 or CB4856 allelic variants of *amx-2* in an *amx-2(lf);let- 60(gf)*. (**A**) Position of the peaks of the eQTL on chromosome I and eQTL N2 allelic effects. Red to blue gradient shows the log2 expression ratio between the allelic variants of *amx-2* in the transgenic lines. QTLs detected in *Schmid et al*. 2015 (QTL1a and QTL1b) and newly identified sub-*trans*-band QTL1c are shown on top. Position of *amx-2* is shown in red. (**B**) Direct comparison of the eQTL effects and transgene effects for all eQTL on chromosome I. (**C**) Direct comparison of the eQTL effects and transgene effects for eQTL in QTL1b/*amx-2* locus. (**D**) Direct comparison of the eQTL effects and transgene effects for eQTL in QTL1c.

**Supplemental table S1:** The genetic map of the *let-60(gf)* sensitized RIL population and the two parental strains used in the gene expression experiment. The CB4856 genotype is denoted with −1 and the N2 genotype is denoted with 1.

**Supplemental table S2:** FLP makers used for this study

**Supplemental table S3:** The assigned gene expression markers, organized per strain per sample per marker. The number of spots on which the correlation was based and the correlation value is given, as well as the assigned genotype.

**Supplemental table S4:** A table summarizing the results of the power analysis. Per simulated peak size (sigma) the variation explained is shown. These peaks were simulated 10 times at each marker location in noise simulated by a standard normal distribution. The number of correctly detected, false-positive, and undetected QTL is shown (at –log10(p) > 3.2). Also the fractions of the total are given. The quantiles of the effect estimation are listed as well as the quantiles of the ‘detected peak’-‘true peak’ distance.

**Supplemental table S5:** A table with the eQTL mapped in the *let-60(gf)* miRIL population. eQTL are listed per trait (Spot) and QTL type. The peak location and its confidence interval are given (based on a 1.5 LOD drop), the peak significance, its effect, and the variation explained by the QTL. The effect direction indicates higher in N2 (positive numbers) or higher at CB4856 (negative numbers) loci. Furthermore, information about the affected gene represented by the microarray spot is shown (name, and location).

**Supplemental table S6:** eQTL enrichments split out by eQTL type. The genotype behind an eQTL class (*e.g*. cis_CB4856) indicates it concerns an eQTL effect with high expression associated with that genotype. The database used for enrichment (Annotation), the category (Group), and the number of genes on the array that are in the group (Genes_in_group) are indicated. Furthermore, the overlap with the cluster (Overlap) and the Bonferroni-corrected significance of that overlap are shown.

## ACKNOWLEDGEMENTS

The authors thank Harm Nijveen for making our data available in EleQTL.

## AUTHOR CONTRIBUTIONS

AH, JK, and LBS conceived and designed the experiments. JAGR, MR, and TS conducted the experiments. MGS, LBP, and LBS conducted transcriptome and main analyses. MGS, AH, JK, and LBS wrote the manuscript.

